# THE FIRST COMPREHENSIVE FORENSIC GENETIC STUDY OF A MEMBER OF CHRISTOPHER COLUMBUS’ FAMILY: HIS GREAT-GREAT-GRANDSON

**DOI:** 10.1101/2024.10.12.618003

**Authors:** I Navarro-Vera, J Yravedra Saez de los Terreros, A Bonilla, M Tirapu, M Albert, P Jímenez, D Herránz, C García

**Affiliations:** Forensic Genetics Department, Citogen, Zaragoza; Universidad Complutense de Madrid; P y A Arqueologos SL; Independent Anthropology Researcher; Mirella Albert, SL; Independent Researcher

**Keywords:** ancient DNA, forensic genetic, MPS, flanking regions, ancestral SNPs, phenotypes, autosomal DNA

## Abstract

The present study has successfully obtained genetic information that allow the identification from direct descendants of Christopher Columbus, dating from 1526 to 1734. This represents the first achievement of this unique objective. The results obtained from the analysed samples, characterized by their significant antiquity, yield exceptional insights into forensic genetics and its newest protocols and methodologies applied to the study of ancient skeletal remains. This work presents the preliminary results of the whole family study using massive parallel sequencing, focusing on the autosomal Single Nucleotide Polymorphism (SNP) and Short Tandem Repeats (STR) markers, including their flanking region sequences, as well as ancestry informative markers (AIMS) of the great-great-grandson of the discoverer.

The results of this project pave the way towards the application of new sequencing technologies for the study of genetic evolution, biogeographical origins and forensic identification of historical characters.

## INTRODUCTION

A family pantheon founded in the 16th century with several buried members of Cristopher Columbus descendants has provided the unique opportunity to conduct this research. These noble direct descendants of Christopher Columbus were buried in a crypt located inside the church of Santa María de Gracia in the municipality of Gelves in the province of Sevilla (Spain). There is wide historical documentation and also oral tradition that confirm both location and family members present in the mentioned burial. The presence of the coat of arms of Christopher Columbus’ family and the Royal House of Portugal also identify this noble family in this place. In addition, the crypt contains three niches with doors and presents the characteristic architecture layout of a Portuguese burial place.

Access to the crypt has been guarded since the 15th century by ecclesiastic authority and remained restricted and closed under lock and key. After obtaining all the necessary permits to access the family pantheon, the archaeology and anthropology team of this research group was able to access and begin its work in the crypt, located under the church. After the archeologic and anthropologic study was made in the crypt, bone samples from the different individuals were collected.

Among all the descendants of Christopher Columbus deposited in the burial, the present study focuses on his great-great-grandson -Jorge Alberto de Portugal, whose initial identification was carried out on the basis of historical documentation, archaeological and anthropological findings, such as age estimation (around twenty years at death time and bone density among other data). His skeletal remains were in an excellent conservation status. To perform the genetic analysis a femur sample was selected and sent to the laboratory to be processed. The genetic study was carried out in an accredited forensic genetic laboratory applying novel molecular techniques such as next generation sequencing. To our knowledge this is the first time where a complete autosomal STR profile, including the study of their flanking regions as well as ancestry and phenotypic SNPs markers have been obtained from such an ancient skeletal remain, and even more important from a direct descendant of Christopher Columbus.

These genetic data will help to understand who Christopher Columbus’ family was, their ancestral biogeographical origins and will undoubtedly contribute to kinship studies and identification of further family members.

## MATERIAL AND METHODS ^**1-12**^

### Bone samples

Several bone fragments were gained from the femur sample obtained in the crypt that belonged to the individual identified as the great-great-grandson of Christofer Columbus.

### DNA lysis, DNA extraction and DNA quantification

Two different extraction procedures from different bone pieces were carried out in simultaneously. After cleaning and disinfection, bone samples were grinded using a Tissuelyser II (QIAGEN). Bone powder was incubated O/N in a lysis/decalcification buffer. Subsequent DNA extraction was carried out using PrepFiler™ BTA (ThermoFischer). DNA concentration process was carried out using Amicon®Ultra-0.5 columns (Millipore). The obtained DNA was quantified by real-time PCR using QuantifilerTM Trio DNA Quantification kit in a QuantStudio5 thermocycler (ThermoFischer).

### Library preparation and targeted sequencing

The extracted DNA from each fragment was analysed in duplicate. For each replicate, 125 pg DNA input were used for library preparation using the Primer Mix B of the ForenSeq™ DNA Signature Prep kit (Verogen) following manufacturer’s protocol. One negative PCR control, one negative DNA extraction control and one positive control (2800M provided in the kit) were included as part of the processed samples. Libraries were then purified and normalized following manufacturers protocol and pooled together (5 µL each). The denaturalized pool was loaded on a MiSeq® FGx Reagent Cartridge. MPS run was set up in a MiSeq®FGx instrument. Targeted sequencing took place on a Standard MiSeq®FGx Flow Cell (QIAGEN).

### Data analysis using the UAS

Data analyses presented here were performed with the Universal Analysis Software (UAS) applying the manufacturer’s default settings.

## RESULTS

After DNA extraction and subsequent concentration, 40 µL of DNA at a concentration of 0,027 ng/µL were obtained. The genetic material was very good conserved and showed a degradation index below 2. In terms of sequencing, run metrics such as cluster density, phasing, prephasing and passing filter values were in the expected range (1119k/mm^2^, 0.174%, 0.017% and 89.98% respectively). Sample replicates showed an average of 300.000 reads each. Positive sequencing control showed 402.000 reads and 100% concordance with the expected results for STRs and SNPs. Both, library preparation negative control and DNA extraction negative control resulted in 0 reads. The Human Sequencing Control (HSC) used as internal control by the UAS was in the expected quality range and intensity.

All the results presented were concordant between all four sample replicates: two bone sample fragments with two replicates each.

A complete autosomal STR profile was obtained, including 27 markers (see table nº 1) confirming that the present study has obtained the most comprehensive DNA profile from one of the direct descendants of Christopher Columbus.

The flanking regions from all the above mentioned STRs were also sequenced (see Supplementary data). The variants detected in these regions will contribute to a highly accurate kinship analysis leading to a more conclusive pedigree building with the rest of the family members found in the crypt. As already shown by Devesse et al^14^ the study of these regions can also give information about the ancestry of the individual. Moreover, a total of 56 ancestry-informative SNPs (aSNPs) has been genotyped. (See table nº 2).

This study also made it possible to determine a complete Y chromosome STR profile with 24 genotyped markers. The obtained Y-chromosome haplotype was crucial for the identification of the skeletal remains as proceeding from Christopher Columbus great-great-grandson, since it links him directly with the paternal linage of the Royal House of Portugal (data to be published).

**Table 1:** Genetic profile based on autosomal STRs from the Great-Great-Grandson of Christographer Colombus.

| Marker | Allele | Marker | Allele |
| --- | --- | --- | --- |
| Amelogenin | X/Y | D10S1248 | 14/17 |
| D1S1656 | 16/17 | TH01 | 9/9.3 |
| TPOX | 8 | vWA | 14/17 |
| D2S441 | 11/15 | D12S391 | 18 |
| D2S1338 | 19/24 | D13S317 | 9/11 |
| D3S1358 | 14/16 | PentaE | 10/13 |
| D4S2408 | 8/10 | D16S539 | 9/12 |
| FGA | 21/23 | D17S1301 | 11/12 |
| D5S818 | 11/12 | D18S51 | 15 |
| CSF1PO | 10/12 | D19S433 | 13 |
| D6S1043 | 11/17 | D20S482 | 13 |
| D7S820 | 10/13 | D21S11 | 29/30 |
| D8S1179 | 12 | PentaD | 13/14 |
| D9S1122 | 12/13 | D22S1045 | 15/16 |

**Table 2:** Biogeographical Ancestry SNPs genotype of the Great-Great-Grandson of Christographer Colombus.

| Biogeographical ancestry markers |  |  |  |  |  |  |  |  |  |
| --- | --- | --- | --- | --- | --- | --- | --- | --- | --- |
| rs3737576 | AA | rs4833103 | CC | rs1871534 | CC | rs200354 | GG | rs12498138 | GG |
| rs7554936 | CT | rs1229984 | AG | rs3814134 | TT | rs1800414 | AA | rs2196051 | TC |
| rs2814778 | AA | rs3811801 | CC | rs4918664 | AG | rs12439433 | AA | rs9522149 | CC |
| rs798443 | GA | rs7657799 | TT | rs174570 | CC | rs735480 | TC | rs7226659 | GG |
| rs1876482 | CC | rs870347 | TT | rs1079597 | GG | rs459920 | CC | rs3916235 | GG |
| rs1834619 | GG | rs7722456 | CT | rs2238151 | TC | rs4411548 | GA | rs4891825 | AA |
| rs3827760 | TT | rs192655 | AA | rs671 | GG | rs2593595 | TT | rs7251928 | AC |
| rs260690 | AA | rs3823159 | AA | rs7997709 | TT | rs17642714 | AA | rs310644 | AA |
| rs6754311 | CC | rs917115 | TC | rs1572018 | GG | rs4471745 | GG | rs2024566 | AG |
| rs10497191 | TC | rs1462906 | CC | rs2166624 | GA | rs11652805 | CT | rs16891982 | GG |
| rs1919550 | AA | rs6990312 | GT | rs7326934 | GG | rs2042762 | AA | rs12913832 | AA |

## DISCUSSION

This study achieved to determine more than 200 forensically relevant genetic markers from the great-great-grandson of Christopher Columbus by applying MPS. 27 autosomal short tandem repeats (aSTRs), 7 X chromosomal STRs (X-STRs), 24 Y chromosomal STRs (Y-STRs), 94 identity-informative single nucleotide polymorphisms (iSNPs), 22 phenotypic-informative SNPs (pSNPs) and 56 ancestry-informative SNPs (aSNPs), were consistently determined.

To our knowledge, this is the first study that allows a scientifical insight into the DNA of Christopher Columbus direct descendants. Classically, one of the mayor difficulties in the genetic study of ancient skeletal remains is the low amount of recovered DNA, that is also usually highly degraded. The application of novel molecular techniques for DNA extraction and sequencing and the fact of the remains being well conserved in a crypt made it possible to get this enormous amount of genetic information out of such an ancient sample.

Moreover, forensic investigative genetic genealogy (FIGG) techniques have been applied to study different members of this burial (data still not published), showing again that the application of novel forensic genetic techniques to ancient bone samples can lead to promising results.

## CONCLUSION

Data shown in this article reveal the most comprehensive DNA profile from a direct descendant of Christopher Columbus achieved to date: Jorge Alberto de Portugal, his great-great-grandson. The genetic markers obtained allowed to scientifically link the skeletal remains to the Royal House of Portugal, to get information about the biogeographical ancestry, phenotype and genetic identification of this historical character.

At this point of time, genetic data on five other family members has been obtained (data still not published). These data are being analysed to obtain scientific evidence and a greater inshight into the genetic background of Christopher Columbus and his family. At the time of submission of this article, a much larger number of SNPs from the Jorge Alberto de Portugal, the great-great-grandson from Columbus are already available and are being studied to establish kinship to other individuals found in the crypt.

The work shows the preliminary phase of a much more ambitious study. The family group found in the crypt is being analysed in order to establish a clear genealogical tree. Genetic forensic genealogy studies (data not shown) have been carried out on different members of the burial. These studies promise to shed light on the true family of Columbus and will be the first step to clarify the historical doubt about the true origins of Christopher Columbus.

This text is pending from revision so some changes in the wording of the text may occur.

## Supporting information

Supplementary table 1: STR Flanking Regions

## ACKNOWLEGEMENTS

To all the members of the “Archidiocésis de Sevilla” with specially thanks to Don Teodoro León Muñoz, bishop and general vicario of the “Archidiócesis de Sevilla”. Thank you so much for the authorisation which made this scientific project possible. And to Alba Hernández for her meticulous and accurate work in the laboratory.

