## Supplementary table 1: STR Flanking Regions for "THE FIRST COMPREHENSIVE FORENSIC GENETIC STUDY OF A MEMBER OF CHRISTOPHER COLUMBUS’ FAMILY: HIS GREAT-GREAT-GRANDSON"

### Autosomal STR Flanking Region Report

| Note | |
|---|---|
| | Sequence column D contains flanking region and repeat region amplicon sequence data exclusive of the ForenSeq PCR primers.<br>Some loci may have partially truncated downstream sequences. (refer to the ForenSeq Universal Analysis Software User Guide).<br>Nucleotide differences present within the target region reported in the ForenSeq Universal Analysis Software (e.g. repeat region) are presented in bold, black text.<br>Nucleotide differences identified outside of the target region (e.g. flanking region) are presented in underlined, bold, blue text. |

**Autosomal STR Coverage Information**

| Locus | STR Length | Read | Sequence | Strand |
|---|---|---|---|---|
| Amelogenin | 0 | 215 | TAGTGTGTTGATTCTTTATCCCAGATGTTTCTCAAGTGGTCCTGATTTTACAGTTCCTACCAC | Forward |
| Amelogenin | 6 | 828 | TAGTGGGTGGATTCTTCATCCCAAATAAAGTGGTTTCTCAAGTGGTCCCAATTTTACAGTTCCTACCAT | Forward |
| D1S1656 | 16 | 442 | TAGATAGATAGATAGATAGATAGATAGATAGATAGATAGATAGATAGATAGATAGATAGATAGATAGGTGTGTGTGTGTTTAATTGTATGTATATATATTTGGTTCCCTAGTGATTCTATTTCTCTGAA | Reverse |
| D1S1656 | 17 | 459 | TAGATAGATAGATAGATAGATAGATAGATAGATAGATAGATAGATAGATAGATAGATAGATAGATAGATAGGTGTGTGTGTGTTTAATTGTATGTATATATATTTGGTTCCCTAGTGATTCTATTTCTCTGAA | Reverse |
| TPOX | 8 | 2025 | AATGAATGAATGAATGAATGAATGAATGAATGTTTGG | Forward |
| D2S441 | 15 | 1080 | CCAGGAACTGTGGCTCATCTATGAAAACTTCTATCTATCTATCTATCTATCTATCTATCTATCTATCTATCTATCTATTTATCTATCTATA**T**CATAACACCACAGCCACTTA | Forward |
| D2S441 | 11 | 1525 | CCAGGAACTGTGGCTCATCTATGAAAACTTCTATCTATCTATCTATCTATCTATCTATCTATCTATCTATCTATATCATAACACCACA**G**CCACTTA | Forward |
| D2S1338 | 24 | 594 | AAATGGCTTGGCCTTGC**C**TGCCTGCCTGCCTGCCTGCCTTCCTTCCTTCCTTCCTTCCTTCCTTCCTTCCTTCCTTCCTTCCTTCCTTCCTTCCTTCCGTCCTTCCTTCCCTC | Reverse |
| D2S1338 | 19 | 1318 | A**A**ATGGCTTGG**C**CTTGCCTGCCTGCCTGCCTGCCTGCCTGCCTTCCTTCCTTCCTTCCTTCCTTCCTTCCTTCCTTCCTTCCTTCCTTCCCTC | Reverse |
| D3S1358 | 14 | 2191 | TTTGGGGGCATCTCTTATACTCATGAAATCAACAGAGGCTTGCATGTATCTATCTGTCT**G**TCT**A**TCTATCTATCTATCTATCTATCTATCTA**T**CTATCTATCTATGAGACAGGGTCTTGCTC | Forward |
| D3S1358 | 16 | 3062 | TTTGGGGGCATCTCTTATACTCATGAAATCAACAGAGGCTTGCATGTATCTATCTGTCTGTCT**G**TCT**A**TCTATCTATCTATCTATCTATCTATCTA**T**CTATCTA**T**CTATCTATCTATGAGACAGGGTCTTGCTC | Forward |
| D4S2408 | 10 | 1425 | CTATGCATC**T**ATCTATCTATCTATCTATCTATCTATCTATCTATCTAATGGTTA | Reverse |
| D4S2408 | 8 | 2251 | CTATGCATCTATCTATCTATCTATCTATCTATCTATCTAATGGTTA | Reverse |
| FGA | 21 | 1513 | G**C**ATATTTACAAGCTAGTTTCTTTCTTTCTTTTTTCTCTTTCTTTCTTTCTTTCTTTCTTTCTTTCTTTCTTTCTTTCTTTCTTTCTTTCTCCTTCCTTCCTTTCTTCCTTTCTTTTTTGCTGG | Reverse |
| FGA | 23 | 2271 | G**C**ATATTTAC**A**AGCTAGTTTCTTTCTTTCTTTTTTCTCTTTCTTTCTTTCTTTCTTTCTTTCTTTCTTTCTTTCTTTCTTTCTTTCTTTCTTTCTTTCTCCTTCCTTCCTTTCTTCCTTTCTTTTTTGCTGG | Reverse |
| D5S818 | 12 | 312 | ATTTTGAAGATAGATAGATAGATAGATAGATAGATAGATAGATAGATAGATAGATAGATAGAGGTATAAATA | Forward |
| D5S818 | 11 | 607 | ATTTTGAAGATAGATAGATAGATAGATAGATAGATAGATAGATAGATAGATAGA**T**GTATAAATA | Forward |
| CSF1PO | 12 | 441 | AAGATAGATAGATTAGATAGATAGATAGATAGATAGATAGATAGATAGATAGATAGATAGATAGGAAG | Forward |
| CSF1PO | 10 | 458 | AAGATAGATAGATTAGATAGATAGATAGATAGATAGATAGATAGATAGATAGATAGGAAG | Forward |
| D6S1043 | 17 | 1189 | GATCAATAGATTGATAGATCAGATAGATAGATAGATAGATAGATAGATAGATAGATAGATAGATAGATACATAGATAGATAGATAGATAGATAGGATTTATTATGGGAAGTGGCTCATACAATTATGGAA**G**CTGAGAAATTTCACAATATGCCATCT | Forward |
| D6S1043 | 11 | 1412 | GATCAATAGATTGATAGATCAGATAGATAGATAGATAGATAGATAGATAGATAGATAGATAGATAGGATTTATTATGGGAAGTGGCTCATACAATTATGGAA**G**CTGAGAAATTTCACAATATGCCATCT | Forward |
| D7S820 | 13 | 414 | ATAGTTTAGAACGAACTAA**C**GATAG**A**TAG**A**TAGATAGATAGATAGATAGATAGATAGATAGATAGATAGATAGACAGATTGATAGTTTTTTTT**TATCTCACT**AA**ATA** | Reverse |
| D7S820 | 10 | 521 | ATAGTTTAGAA**C**GAACTAACGATAGATAGATAGATAGATAGATAGATAGATAGATAGATAGACAGATTGATAGTTTTTTTTTATCTCACTAAATA | Reverse |
| D8S1179 | 12 | 1686 | TCTATCTATCTATCTATCTATCTATCTATCTATCTATCTATCTATCTATTCCC | Forward |
| D9S1122 | 12 | 1754 | AGATAACTGTAGATAGGTAGAT**A**GATAGATAGATAGATAGATAGATAGATAGATAGATAGATAGATATTAAT | Reverse |
| D9S1122 | 13 | 2047 | AGATAACTGTAGATAGGTAGAT**C**GATAGATAGATAGATAGATAGATAGATAGATAGATAGATAGATAGATATTAAT | Reverse |
| D10S1248 | 17 | 918 | TTGAACAAATGAGTGAGTGGAAGGAAGGAAGGAAGGAAGGAAGGAAGGAAGGAAGGAAGGAAGGAAGGAAGGAAGGAAGGAAGGAAGGAAAT**G**AA**G**ACAATACAACCAGAGTT | Forward |
| D10S1248 | 14 | 1245 | TTGAACAAATGAGTGAGTGGAAGGAAGGAAGGAAGGAAGGAAGGAAGGAAGGAAGGAAGGAAGGAAGGAAGGAAATGAAGACAA**T**ACAA**CC**AGAGT**T** | Forward |
| TH01 | 9 | 4073 | TGCAGGTCACAGGGAACACAGACT**C**CATGGTGAATGAATGAATGAATGAATGAATGAATGAATGAATGA**G**GGAAATAAGG | Reverse |
| TH01 | 9.3 | 5410 | TG**C**AGGT**C**ACAGGGAA**C**ACAGACT**C**CATGGTG**A**ATGAATGAATGAATGAATGAATGATGAATGAATGAATGAGGGAAATAAGG | Reverse |
| vWA | 17 | 157 | ATTGATCTATCTGTCTGTCTGTCTGTCTGTCTATCTATCTATCTATCTATCTATCTATCTATCTATCTATCTATCTATCCATCTATCCATCCATCCTATGTATT | Forward |
| vWA | 14 | 462 | ATTGATCTATCTGTCTGTCTGTCTGTCTATCTATCTATCTATCTATCTATCTATCTATCTATCCATCTATCCATCCATCCTATGTATT | Forward |
| D12S391 | 18 | 681 | CAGAGAGAAAGAATCAACAGGATCAATGGATGCATAGGTAGATAGATAGATAGATAGATAGATAGATAGATAGATAGATAGATAGA**T**AGA**C**AGACAGACAGACAGACAGACAGATGAGAGGGGATTTATTAGAGGAATTAGCTCAAGTGATATGGAGGCTGAAAAATCT**C**ATGACAGTCCATCTGCAAGCT**G**GA**G**ACCCAG**G**GACACTA | Forward |
| D13S317 | 11 | 553 | TCTGACCCATCTAACGCCTATCTGTATTTACAAATACATTATCTATCTATCTATCTATCTATCTATCTATCTATCTATCTATCAATCATCTATCTATCTTTCTGTCTGTCTTTTTGGG | Reverse |
| D13S317 | 9 | 1328 | TCTGACCCATCTA**A**CGCCTATCTGTATTTACAAATACATTATCTATCTATCTATCTATCTATCTATCTATCTATCAATCAATCATCTATCTATCTTTCTGTCTGTCTTTTTGGG | Reverse |
| PentaE | 13 | 120 | AAAGAAAAGAAAAGAAAAGAAAAGAAAAGAAAAGAAAAGAAAAGAAAAGAAAAGAAAAGAAAAGAAAATTGTAAGGAGTTTTCT | Reverse |
| D16S539 | 9 | 883 | TCCTCTTCCCTAGATCAATACAGACAGACAGACAGGTGGATAGATAGATAGATAGATAGATAGATAGATAGATATCATTGAAAGACAAAACAGAGATGGA**T**GATAGATAC | Forward |
| D16S539 | 12 | 1424 | TCCTCTTCCCTAGATCAATACAGACAGACAGACAGGTGGATAGATAGATAGATAGATAGATAGATAGATAGATAGATAGATAGATATCATTGAAA**G**ACAAAA**C**AGA**G**ATGGA**T**GA**T**AGATAC | Forward |
| D17S1301 | 11 | 2733 | ATATGTGTGAGATAGATAGATAGATAGATAGATAGATAGATAGATAGATAGATAGATCCATCATAGGAATTTT | Forward |
| D17S1301 | 12 | 3373 | ATATGTGTGAGATAGATAGATAGATAGATAGATAGATAGATAGATAGATAGATAGATAGATCCATCATAGGAATTTT | Forward |
| D18S51 | 15 | 1997 | GTCTCAGAAAGAAAGAAAGAAAGAAAGAAAGAAAGAAAGAAAGAAAGAAAGAAAGAAAGAAAGAAAGAAAAAGAGAGAGGAAAGAAAGAGAAAAAGAAAAGAAATAGTAGCAACTG**T**TATTGTAAGA | Forward |
| D19S433 | 13 | 926 | AGCTATAATTGT**A**CCACTGCACTCCAGCCTGGGCAACAGAATAAGATTCT**G**TTGAAGGAAAGAAGGTAGGAAGGAAGGAAGGAAGGAAGGAAGGAAGGAA**G**GAAGGAAGGAGAGAGGAAGAAAGAGAGAAGATTTTTATT | Reverse |
| D20S482 | 13 | 7979 | AGACACCGAACCAATAAGAGATAGATAGATAGATAGATAGATAGATAGATAGATAGATAGATAGATAGATAGATAGAGATTTATTATAG**G**AA**TTG**A**TT** | Forward |
| D21S11 | 29 | 361 | AAATATGTGAGTCAATTCCCCAAGTGAATTGCCTTCTATCTATCTATCTATCT**G**TCT**G**TCTGTCTGTCTGTCTGTCT**A**TCTATCTAT**A**T**C**TATCTATCTATC**ATCT**ATCTATC**C**AT**ATC**TATCTATCTATCTATCTATCTATCTATCTATCTATCTATC**G**TCTATCTAT | Forward |
| D21S11 | 30 | 689 | AAATATGTGAGTCAATTCCCCAAGTGAATTGCCTTCTATCTATCTATCTATCTATCT**A**TCTGTCTGTCTGTCTGTCTGTCTATCTATCTATATCTATCTATCTATCATCTATCTATCCATATCTATCTATCTATCTATCTATCTATCTATCTATCTATCTATC**T**ATC**G**TCTATCTAT | Forward |
| PentaD | 13 | 59 | GATCACTTGAGCCTGGAAGGTCGAAGCTGAAGTGAGCCATGATCACACCACTACACTCCAGCCTAGGTGACAGAGCAAGACACCATCTCAAGAAAGAAAAAAAAGAAAGAAAAGAAAAGAAAAGAAAAGAAAAGAAAAGAAAAGAAAAGAAAAGAAAAGAAAAGAAAAGAAAAGAAAAGAAAAAACGAAG**GGG**AAAAAAAGAGAATCATAAACATAAATGTAAAATTTCTCAA | Forward |
| PentaD | 14 | 93 | GATCACTTGAGCCTGGAAGGTCGAAGCTGAAGTGAGCCATGATCACACCACTACACTCCAGCCTAGGTGACAGAGCAAGACACCATCTCAAGAAAGAAAAAAAAGAAAGAAAAGAAAAGAAAAGAAAAGAAAAGAAAAGAAAAGAAAAGAAAAGAAAAGAAAAGAAAAGAAAAGAAAAGAAAAGAAAAAAC**G**AAG**G**G**G**AAAAAAAGAGAATCATAAACATAAATGTAAAATTTCT | Forward |
| D22S1045 | 16 | 971 | CGTT**G**GAATTC**C**CCAAACTGG**C**CAGTTCCTCTCCACCCTATAGACCCTGTCCTAGCCTTCTTATAGCTGCTATGG**G**GGCTAGATTTTCCCCGATGATAGTAGTCTCATTATTATTATTATTATTATTATTATTATTATTATTACTATTATTGTTATAAAAATATTGCCAAT | Reverse |
| D22S1045 | 15 | 1520 | C**G**TTGGAATT**CC**CCAAACTGGCCAGTTCCTCTCCACCCTATAGACCCTGTCCTAGCCTTCTTATAGCTGCTATGGGGGCTAGATTTTCCCCGATGATAGTAGTCTCATTATTATTATTATTATTATTATTATTATTATTACTATTATTGTTATAAAAATATTGCCAAT | Reverse |
